# DNA Damage Accelerates G-Quadruplex Folding in a Duplex-G-Quadruplex-Duplex Context

**DOI:** 10.1101/2024.01.20.576387

**Authors:** Aaron M. Fleming, Brandon Leonel Guerra Castañaza Jenkins, Bethany A. Buck, Cynthia J. Burrows

## Abstract

Molecular details for DNA damage impact on the folding of potential G-quadruplex sequences (PQS) to non-canonical DNA structures that are involved in gene regulation are poorly understood. Here, the effects of DNA base damage and strand breaks on PQS folding kinetics were studied in the context of the *VEGF* promoter sequence embedded between two DNA duplex anchors, referred to as a duplex-G-quadruplex-duplex (DGD) motif. This DGD scaffold imposes constraints on the PQS folding process that more closely mimic those found in genomic DNA. Folding kinetics were monitored by circular dichroism (CD) to find folding half-lives ranging from 2 s to 12 min depending on the DNA damage type and sequence position. The presence of Mg^2+^ ions and the G-quadruplex (G4)-binding protein APE1 facilitated the folding reactions. A strand break placing all four G runs required for G4 formation on one side of the break accelerated the folding rate by >150-fold compared to the undamaged sequence. Combined 1D ^1^H-NMR and CD analyses confirmed that isothermal folding of the *VEGF*-DGD constructs yielded spectral signatures that suggest formation of G4 motifs, and demonstrated a folding dependency with the nature and location of DNA damage. Importantly, the PQS folding half-lives measured are relevant to replication, transcription, and DNA repair time frames.

**TOC Graphic:** 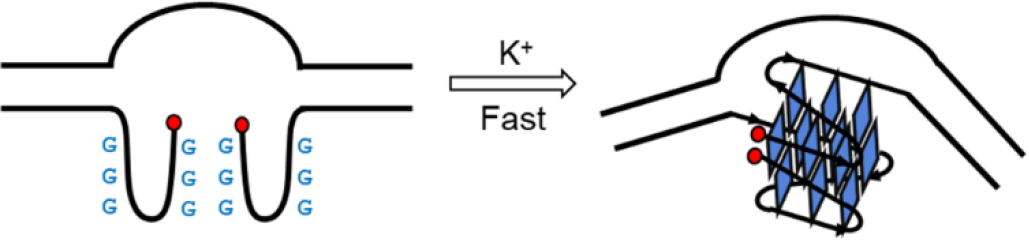

## Manuscript

Here we demonstrate 2’-deoxyguanosine (G) nucleotide DNA damage accelerates a G-quadruplex (G4)-forming sequence folding rate in a duplex-G-quadruplex-duplex (DGD) context. Cellular oxidative stress necessitates transcriptional level changes that can be driven by two properties of G-rich DNA:^1^ (1) G-runs are enriched in human gene promoters that display a greater potential G4 sequence (PQS) probability,^2^ and (2) these sequences are more prone to cellular oxidative damage yielding 8-oxoguanine (OG).^3,4^ Short-patch base excision repair (BER) is a multi-step pathway that removes modified bases via a glycosylase to yield an abasic site (AP), which is cleaved by an endonuclease to generate a DNA nick (Figure 1A).^5^ The nick is cleared via a lyase reaction generating a gap, affording polymerase installation of the correct dNTP, followed by ligation to seal the strand (Figure 1A).^5^ Each step in the repair process furnishes a known type of DNA damage structure.^6^ Our prior cell studies identified that the AP-endonuclease APE1 functions on AP sites derived from OG removal in promoters containing the *VEGF* or other PQSs to induce transcription.^1,7^ Indeed, APE1 has been determined to also function as a transcriptional regulatory protein.^8^ Biophysical studies further revealed APE1 tightly binds promoter G4 folds, supporting a role for this non-canonical DNA structure in gene regulation.^9^ Nevertheless, it remains unclear at what point G4 formation is induced during the DNA repair process. Our findings within a more physiological DGD model provide broader insight into the impact of the DNA damage response pathway on G4 folding in the genomic context.

**Figure 1.**
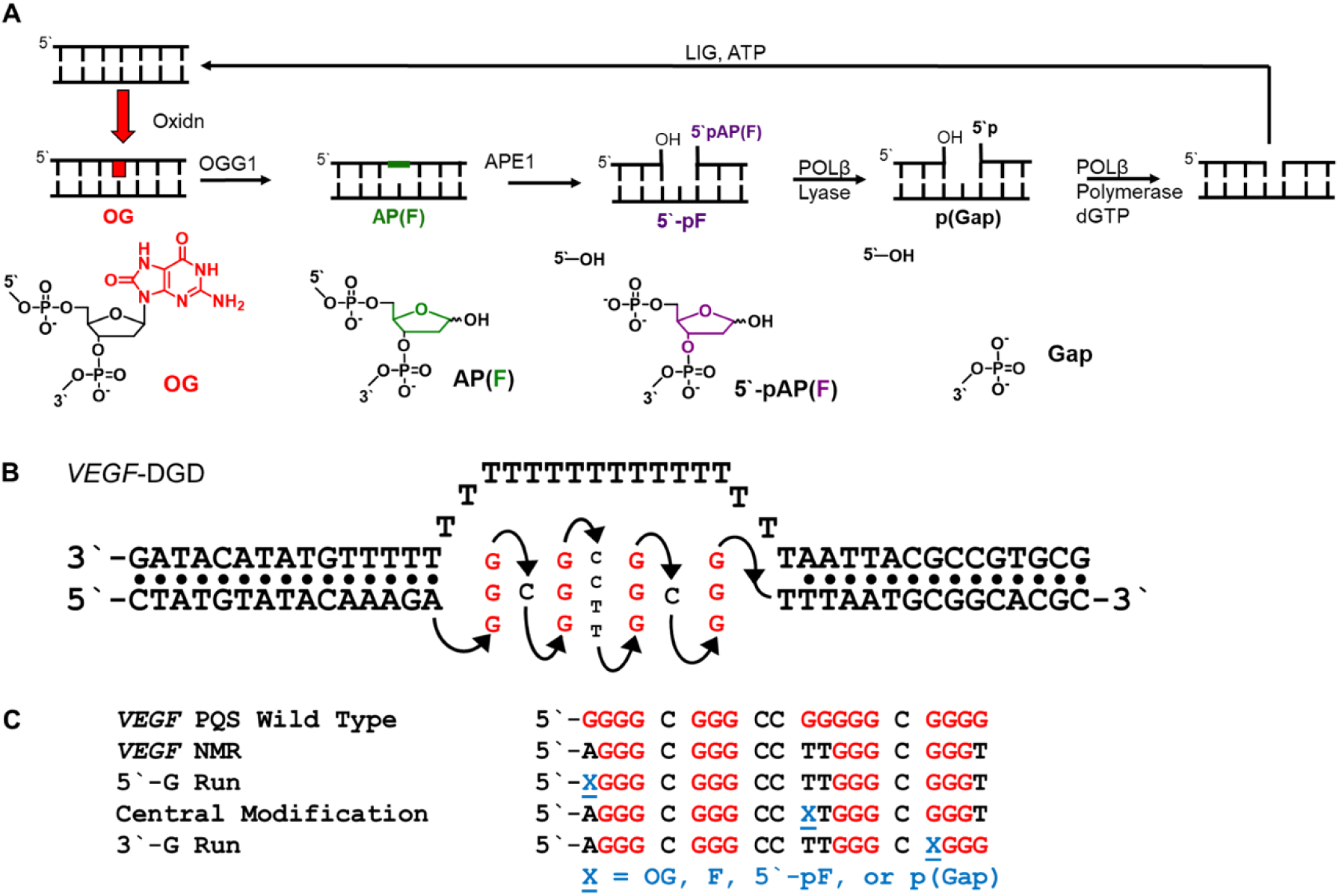
(A) Short-patch BER pathway illustrating the DNA damage structures studied within the *VEGF*-DGD. (B) The DGD construct along with (C) the modified *VEGF* inserts studied in this work.

Inspired by Chaires’ and Trent’s cryo-EM structure of the *c-MYC* G4 embedded between two duplexes (*c-MYC*-DGD),^10^ this DNA framework was used to install the *VEGF* PQS and study its folding (Figure 1B). A critical feature of the *c-MYC*-DGD retained in these studies, was that the complementary strand was modeled with a poly-T segment to connect the flanking DNA duplexes. Specifically, herein the *c-MYC*-DGD sequence was replaced with a *VEGF* sequence that had been previously modified for solution NMR structural investigations in Yang’s laboratory (Figure 1C).^11^ Notably, both the *c-MYC* and *VEGF* promoter G4s adopt parallel-stranded topologies that are dominant folds found in promoter sequences.^12^

A combination of circular dichroism (CD) and 1D ^1^H-NMR analyses were first utilized to verify formation of a *VEGF*-G4 within the DGD scaffold utilizing the previously described thermodynamic folding procedure, which first forms the PQS G4 in the presence of buffered K^+^ ion (140 mM), followed by secondary formation of the duplex flanks through complement strand annealing.^10^ CD analysis of the isolated *VEGF*-G4 produced a spectrum with a 263 nm maximum and 245 nm minimum, while the *VEGF*-DGD showed a 271 nm maximum and 245 nm minimum (Figure 2A). While both spectra are indicative of parallel-stranded G4 formation, the *VEGF*-DGD 8 nm maximum peak redshift is consistent with inclusion of the poly-T complement and duplex handles, which mirrors the previous *c-MYC*-DGD CD spectrum (Figure S2).^10^ In addition, as was observed for the *c-MYC*-DGD, 1D ^1^H-NMR analysis of the *VEGF*-DGD identified two groups of resonance signatures corresponding to the G-H1 and T-H3 imino protons of the duplex DNA (12-14 ppm), and G-H1 iminos involved in Hoogsteen forming G-tetrads (10-12 ppm; Figures 2B and S2).^13^ These initial studies demonstrate that the thermodynamic annealing process affords G4 formation of a *VEGF* PQS within the DGD context.

**Figure 2.**
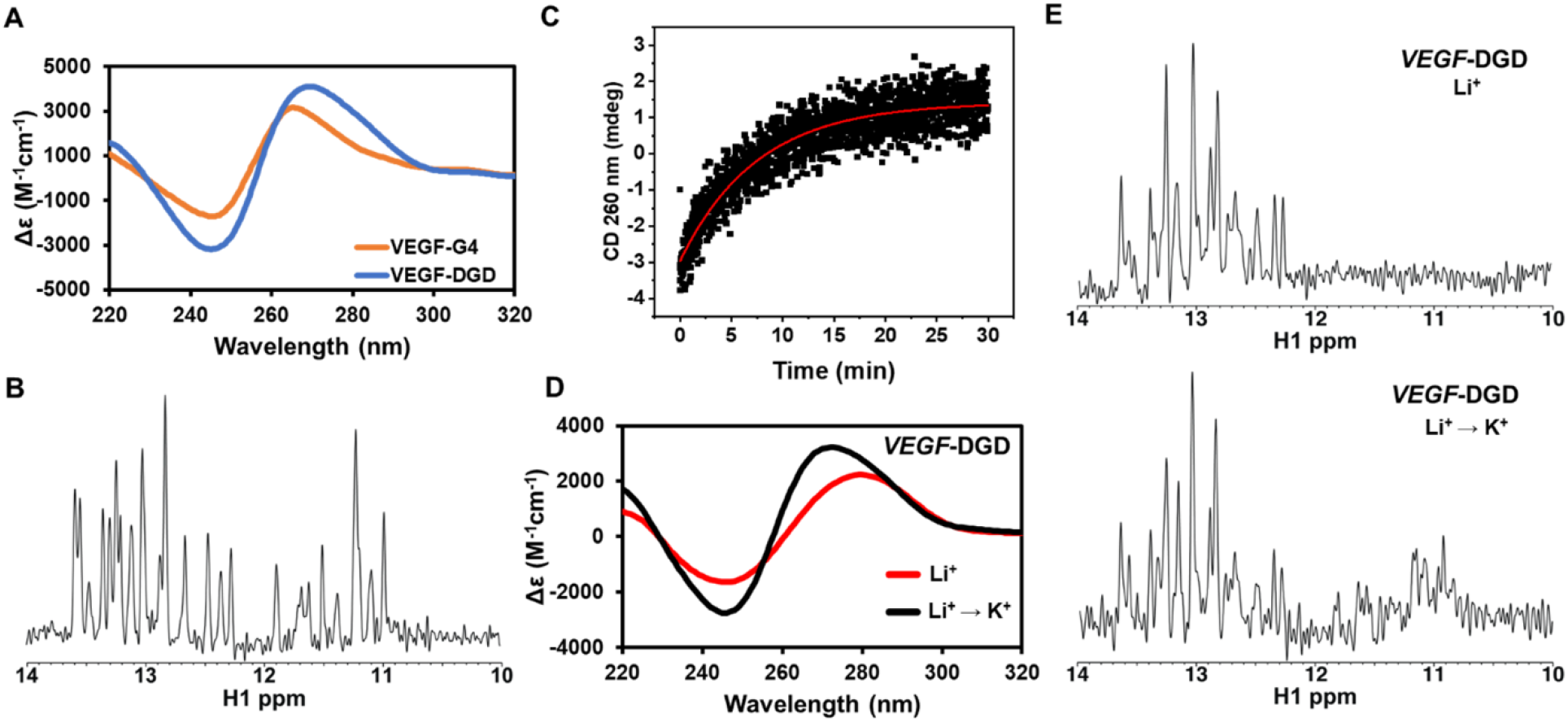
Thermodynamically folded *VEGF*-DGD evaluated by CD (A) and 1D ^1^H-NMR (B). Monitoring of PQS isothermal folding in the DGD context in Li^+^ when 100 mM K^+^ was added by following the CD (C and D) and 1D ^1^H-NMR (E) spectral changes.

Next, we addressed whether the PQS could fold isothermally when embedded between two duplex anchors, which is a situation more analogous to G4 folding in the genome. To generate these constructs, a buffered Li^+^ salt solution was used to first induce formation of the duplex DNA flanks while being agnostic to G4 folding, which enabled a subsequent G4 folding process after addition of K^+^ ions. The duplex arms in 140 mM LiCl had *T*_*m*_ values of 39 and 52 °C, which are consistent with these duplexes being A:T rich (Figure S3). Consequently, the temperature for following folding experiments was held at 30 °C, a value less than the measured *T*_*m*_ values to minimize duplex fraying that would impact G4 folding rate. G4 folding was monitored by time-dependent CD analysis at 260 nm upon addition of 100 mM K^+^, as signal changes at this wavelength directly report on the conversion of an unstructured PQS to a parallel-stranded G4-like fold (Figures 2C/2D).

For the *VEGF*-DGD construct, time evolution of the 260 nm CD signal was hyperbolic and fit to a first-order exponential equation (Figure 2C), as previously described.^14^ The observed rate (*k*_obs_) was 0.13 ± 0.1 min^-1^ corresponding to a folding half-life of 5.3 ± 0.5 min (Figures 3 and S4). The CD spectra before and after K^+^ addition to the Li^+^ sample support folding of a parallel-stranded G4 in this context by the observed changes at 245 and 260 nm (Figure 2D). The spectrum for the isothermally folded sample mirrors that of the thermodynamically folded sample with a lower intensity (Figure 2D). Consistent with CD, the Li^+^ ion sample only exhibited duplex DNA imino peaks by 1D ^1^H-NMR, but after K^+^ addition G:G base pair imino peaks were additionally observed (Figure 2E). Spectroscopic differences between the thermodynamic vs. isothermally folded samples may reflect different final structures or incomplete folding under the timeframe of the analysis, which is not impacted by the presence of Li^+^ ions interfering with the extent of folding as previously reported.^15^ Nonetheless, CD and 1D ^1^H-NMR studies demonstrate the PQS anchored with 5’- and 3’-duplex DNA adopts a G4-consistent fold when K^+^ ions are added. Notably, we determined no significant changes for the rate of PQS folding for poly-T tracks ranging from 14-20 nucleotides, indicating the complement length has minimal influence on the folding reaction (Figure S5).

**Figure 3.**
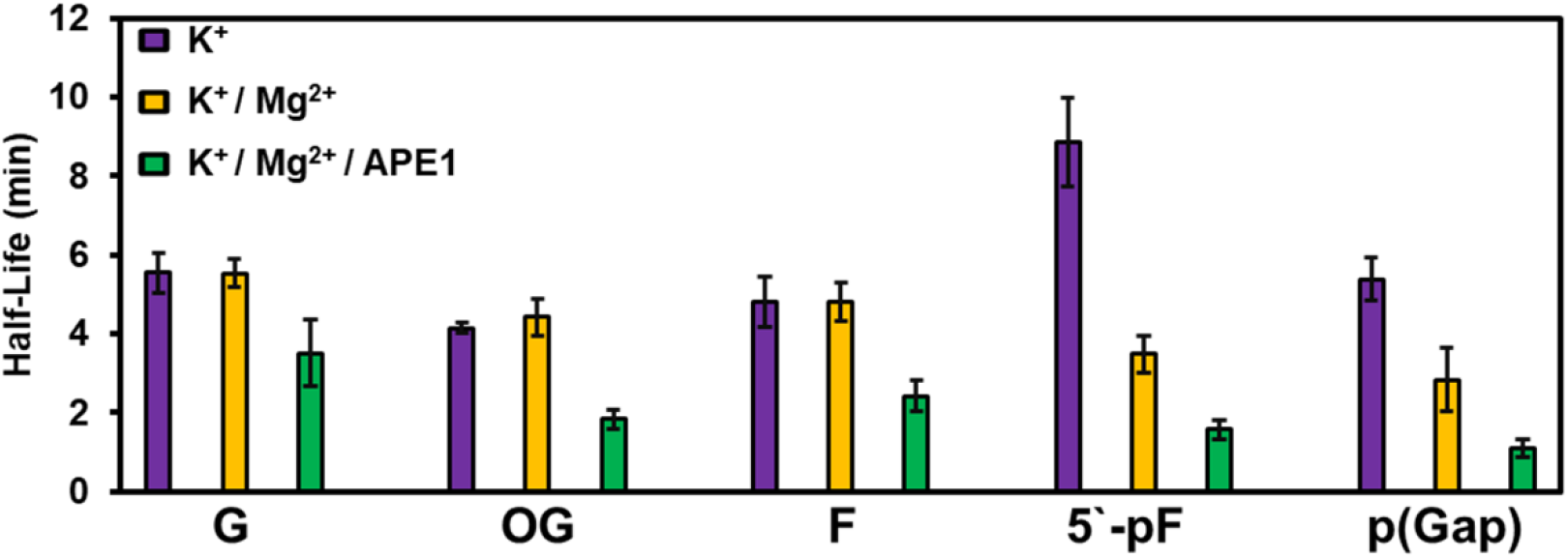
Folding half-lives for the *VEGF*-DGD construct with a central DNA damage when incubated in Li^+^ at 30 °C followed by addition of 100 mM K^+^ ions.

To investigate DNA damage impact on PQS folding within the *VEGF*-DGD scaffold, the PQS-containing strand was synthesized with each stable structure generated during short-patch BER removal of OG to repair it back to G (Figure 1A).^5^ The AP and 5′-pAP structures are chemically unstable, and were synthesized as stable tetrahydrofuran analogs (F and 5′-pF; Figure 1A). Lesions were synthesized at a central G position localized within the loop of the folded G4 that does not impact the core structure, but was previously found to be sensitive to one-electron oxidation, yielding OG (Figure 1C).^16^ Measured folding half-lives for the central OG- and F-containing DGD motifs were similar to the undamaged sequence (4-5 min), while folding for the strand breaks 5′-pF and p(Gap) were slower than or similar to the undamaged sequence, respectively (8 vs. 5 min; Figure 3 and S4). Next, folding studies were conducted with 10 mM Mg^2+^ to determine whether the inclusion of the divalent cation altered folding half-lives, particularly for the 5′ phosphate strand breaks. The only folding half-lives sensitive to Mg^2+^ were the strand breaks, and these decreased by >5-fold to ∼1 min each (Figures 3 and S4). Inspection of the damage-containing DGD sequences in Li^+^ before and after K^+^ addition via CD and 1D ^1^H-NMR support a G4-like motif is formed in each case (Figures 4A-4D and S6). Indeed, 1D ^1^H-NMR spectra of the folded strand break-containing *VEGF*-DGD motifs produced 11 (p(Gap)) or more (5′-pF) G:G base pair imino resonances, strongly supporting the presence of a three G-tetrad G4.

**Figure 4.**
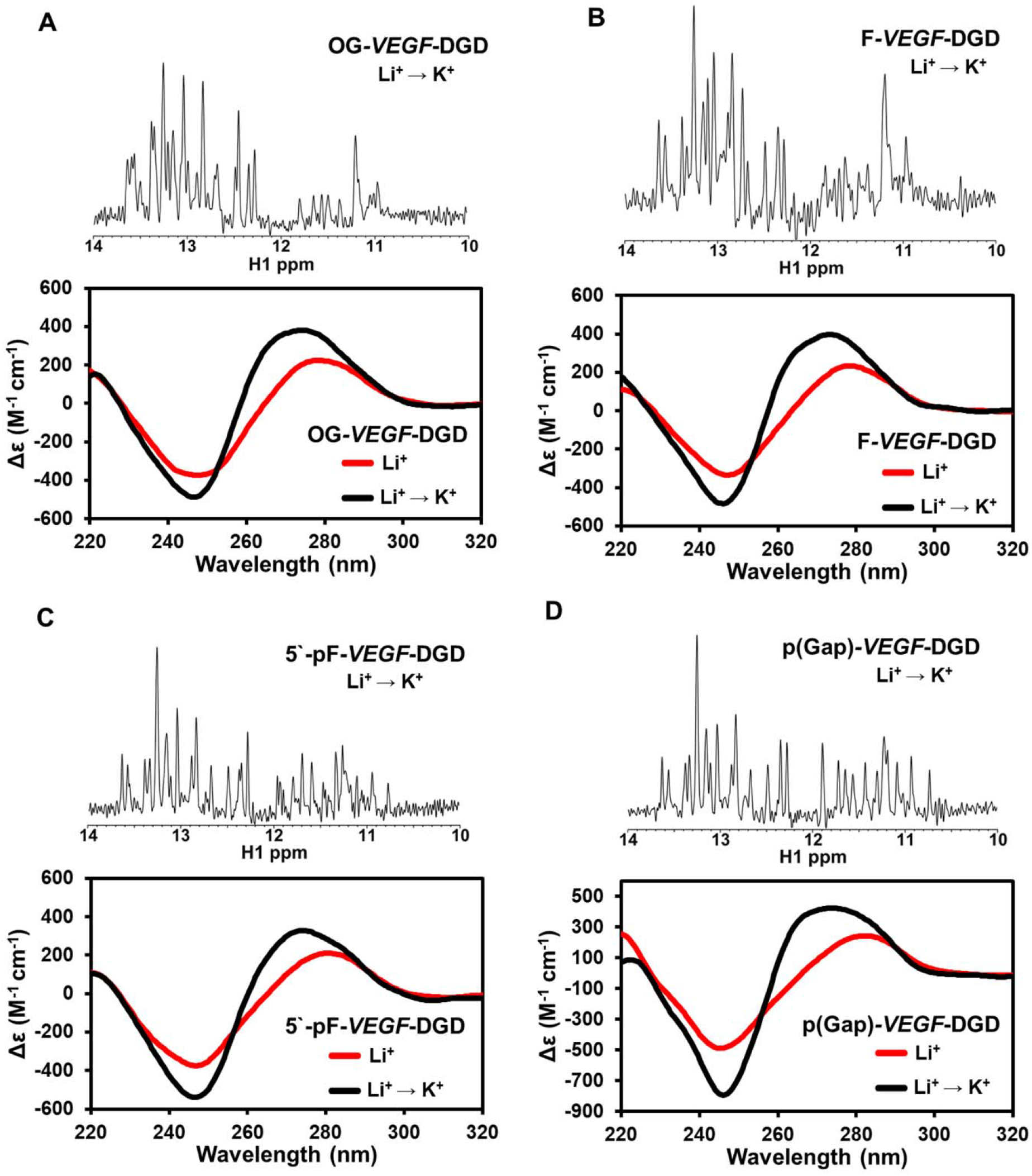
1D ^1^H-NMR and CD characterization of the *VEGF*-DGD motifs containing the DNA damage types (A) OG, (B) F, (C) 5′-pF, or (D) p(Gap).

In our gene activation during oxidative stress proposal, APE1 was required for mRNA induction for *VEGF* promoter PQSs containing OG.^7^ Thus, we investigated whether the inclusion of APE1 during the *VEGF*-DGD folding process, with and without DNA damage, impacted the measured half-life values. Previously, we found APE1 could bind the *VEGF* G4 with or without an F with dissociation constants (*K*_*d*_) around 50 nM.^9^ Similarly, using fluorescence anisotropy, APE1 binding to the *VEGF*-DGD with or without an F had *K*_*d*_ values of ∼50 nM (Figure S7). These binding measurements suggest that APE1 has a strong preference for G4 motifs,^9,17^ and may facilitate PQS folding in the DGD context. When an equimolar amount of APE1 was added to the Li^+^-containing *VEGF*-DGD constructs with K^+^ and Mg^2+^ ions, the folding was consistently faster, providing the lowest folding half-lives (Figures 3 and S4). The largest decrease was for lesion-containing DGD sequences (OG- and F-*VEGF*-DGD ∼2x; 5′-pF- and p(Gap)-*VEGF*-DGD ∼5x; Figure 3). The CD spectra before and after APE1 folding suggest G4-like motifs were formed (Figure S8). Importantly, these studies establish a known G4-binding protein can facilitate folding of a PQS embedded between two duplexes.

The *VEGF* PQS has other sites for which G oxidation was previously observed;^16^ thus, *VEGF*-DGD motifs were designed to place the damage at a known reactive 5′ G on either the 5′-G run or the 3′-G run (Figure 1C). A critical point regarding these positional studies concerns the strand breaks when compared to the studies at the central position. The central strand break will yield strands with two G-runs each that fold together to yield a G4. In contrast, the 5′-G run strand break generates one strand with all four G-runs, and the 3′-G run strand break furnishes two strands that have 3 + 1 G-runs, respectively. Regarding OG and F, we found positional dependency in the folding half-lives, but they were all >2 min with or without Mg^2+^ present (Figures 5 and S9). The strand breaks, 5′-pF and p(Gap), also displayed positional dependency in the folding half-life values in which those on the 3′-G run either did not fold or folded with a half-life value of ∼10 min. In contrast, 5′-G run strand breaks folded with half-life values of <6 s. The 5′-p(Gap) folded fastest with Mg^2+^ present, giving an ∼2 s folding half-life.

**Figure 5.**
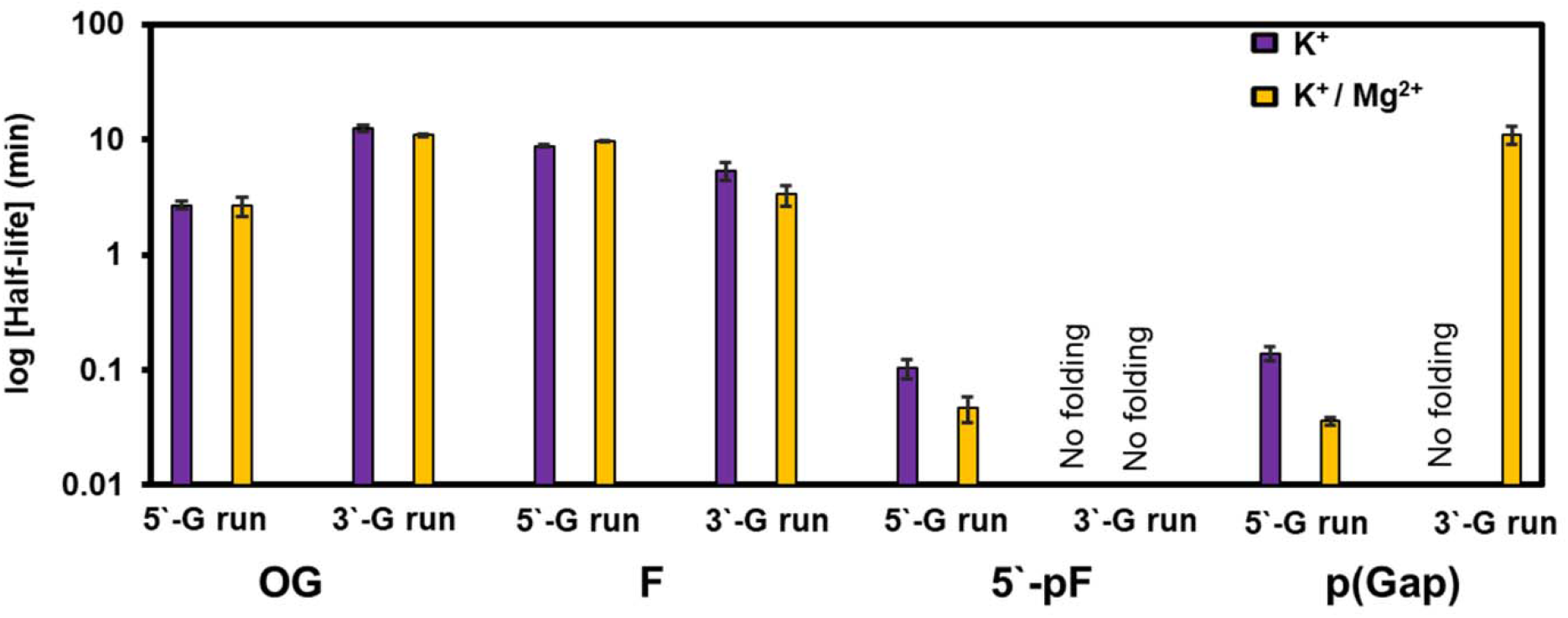
Position-dependent folding half-lives for DNA-damage placed in either 5′-G or 3′-G runs of the *VEGF*-DGD motif.

Compared to the undamaged *VEGF*-DGD, the 5′-p(Gap) folding rate in the 5′-G run was >150-fold faster (Figure 5). CD evaluation before and after folding supports a conclusion that the sequences adopt G4-like motifs in all cases where folding was observed (Figures S9).

Herein, we demonstrate that a PQS can fold when embedded between two duplexes with a model complementary strand analogous to genomic folding. Folding rates measured for the *VEGF*-DGD systems show dependency on DNA damage type and position, as well as the presence of Mg^2+^ ions and a G4-binding protein (Figure 3). The *VEGF* PQS with or without DNA damage in the DGD construct folded with half-lives spanning from 2 s to >10 min. The fastest folding occurs when a strand break places all four G-runs on one side of the break yielding an enhancement of >150-fold compared to the undamaged-DGD system (Figure 5).

Previously, PQS folding was monitored outside the influence and steric constraints of the duplexes at 25 °C yielding folding half-lives of tens of milliseconds,^18,19^ which is ∼200-fold faster than the fastest folder measured herein. We would expect our observed folding rates to increase at 37 °C, though measurements at that temperature would require redesigning the DGD duplex anchors. Nevertheless, the slower observed folding rates for the *VEGF* PQS in the duplex context does not preclude its relevance to biological timeframes. During transcription, R-loops can persist for many minutes,^20^ which is sufficient for at least half of nearly all the *VEGF*-DGD PQSs to fold based on the folding half-lives measured (Figures 3 and 5). Proteins such as APE1 that assist with PQS folding, and are part of DNA repair complexes,^21^ may facilitate folding and expand the structural transition timeframe due to genomic DNA repair initiation taking ∼15 min.^22^ Other G4-interacting proteins may assist in PQS folding,^23^ similar to that observed with APE1. Stalled replication forks can persist for hours, particularly when a PQS is present,^24^ which provides time for these structures to form. Once folded, G4s can serve as hubs for transcription factor binding to impact transcription.^25^

The *VEGF*-DGD system studied does not adequately model the C-rich complementary strand in the folding process. In the cellular context, this strand could be engaged in an R-loop with a G-rich RNA strand or adopt an i-motif structure,^20,26^ the latter being a pH-dependent structure with some folding under neutral cellular conditions.^27^ Future studies to address how R-loops and i-motifs impact PQS folding in the DGD context are needed. Additionally, high-resolution structural analysis of the *VEGF*-DGD constructs with and without DNA damage is required to understand how the duplex anchors and DNA lesions impact G4 structure. Structural studies of the isothermally folded constructs are most important to elucidate whether the topologies generated under more genome-like conditions are similar to thermodynamically folded structures achieved after heating to ∼90 °C followed by cooling. The present results suggest the structures adopted during isothermal folding are G4s, but they do not unequivocally rule out G-triplexes, G-hairpins, or changes in the overall loop and core topologies. We demonstrate a PQS can fold to a non-canonical structure when embedded between duplex DNA segments, and that DNA repair intermediates accelerate the process, which has interesting implications for triggering changes in gene expression.

## Supporting Information

The Supporting Information is available free of charge at https://pubs.acs.org/

Experimental methods, DNA analysis, CD spectra, 1D ^1^H-NMR spectra, *T*_*m*_ analysis, and APE1 binding analysis.

## Acknowledgments

The research was supported by the National Institute of General Medical Sciences (R35 GM145237; C.J.B.), and the University of Utah (B.A.B.). NMR spectra were recorded in the David M. Grant NMR Center, a University of Utah Core Facility. Funds for construction of the NMR Center and the helium recovery system were obtained from the University of Utah and the National Institutes of Health awards 1C06RR017539-01A1 and 3R01GM063540-17W1, respectively. NMR instruments were purchased with support of the University of Utah and the National Institutes of Health award 1S10OD25241-01.

## Notes

### Competing Interest Statement

The authors have declared no competing interest.

## References

1. Fleming, A. M.; Burrows, C. J., Interplay of guanine oxidation and G-quadruplex folding in gene promoters. J. Am. Chem. Soc. 2020, 142, 1115–1136.

2. Chambers, V. S.; Marsico, G.; Boutell, J. M.; Di Antonio, M.; Smith, G. P.; Balasubramanian, S., High-throughput sequencing of DNA G-quadruplex structures in the human genome. Nat. Biotechnol. 2015, 33, 877–881.

3. Ding, Y.; Fleming, A. M.; Burrows, C. J., Sequencing the mouse genome for the oxidatively modified base 8-oxo-7,8-dihydroguanine by OG-Seq. J. Am. Chem. Soc. 2017, 139, 2569–2572.

4. Wu, J.; McKeague, M.; Sturla, S. J., Nucleotide-resolution genome-wide mapping of oxidative DNA damage by click-code-seq. J. Am. Chem. Soc. 2018, 140, 9783–9787.

5. David, S. S.; O’Shea, V. L.; Kundu, S., Base-excision repair of oxidative DNA damage. Nature 2007, 447, 941–950.

6. Gates, K., An overview of chemical processes that damage cellular DNA: spontaneous hydrolysis, alkylation, and reactions with radicals. Chem. Res. Toxicol. 2009, 22, 1747–1760.

7. Fleming, A. M.; Ding, Y.; Burrows, C. J., Oxidative DNA damage is epigenetic by regulating gene transcription via base excision repair. Proc. Natl. Acad. Sci. U.S.A. 2017, 114, 2604–2609.

8. Tell, G.; Quadrifoglio, F.; Tiribelli, C.; Kelley, M. R., The many functions of APE1/Ref-1: not only a DNA repair enzyme. Antioxid. Redox Signal 2009, 11, 601–620.

9. Fleming, A. M.; Howpay Manage, S. A.; Burrows, C. J., Binding of AP endonuclease-1 to G-quadruplex DNA depends on the N-terminal domain, Mg2+, and ionic strength. ACS Bio. & Med. Chem. Au 2021, 1, 44–56.

10. Monsen, R. C.; Chua, E. Y. D.; Hopkins, J. B.; Chaires, J. B.; Trent, J. O., Structure of a 28.5 kDa duplex-embedded G-quadruplex system resolved to 7.4 Å resolution with cryo-EM. Nucleic Acids Res. 2023, 51, 1943–1959.

11. Agrawal, P.; Hatzakis, E.; Guo, K.; Carver, M.; Yang, D., Solution structure of the major G-quadruplex formed in the human VEGF promoter in K+: insights into loop interactions of the parallel G-quadruplexes. Nucleic Acids Res. 2013, 41, 10584–10592.

12. Chen, L.; Dickerhoff, J.; Sakai, S.; Yang, D., DNA G-quadruplex in human telomeres and oncogene promoters: structures, functions, and small molecule targeting. Acc. Chem. Res. 2022, 55, 2628–2646.

13. Plavec, J., NMR Study on Nucleic Acids. In Handbook of Chemical Biology of Nucleic Acids, Sugimoto, N., Ed. Springer Nature Singapore: Singapore, 2022; pp 1–44.

14. Chen, C.; Li, M.; Xing, Y.; Li, Y.; Joedecke, C. C.; Jin, J.; Yang, Z.; Liu, D., Study of pH-induced folding and unfolding kinetics of the DNA i-motif by stopped-flow circular dichroism. Langmuir 2012, 28, 17743–17748.

15. You, J.; Li, H.; Lu, X. M.; Li, W.; Wang, P. Y.; Dou, S. X.; Xi, X. G., Effects of monovalent cations on folding kinetics of G-quadruplexes. Biosci. Rep. 2017, 37, BSR20170771.

16. Fleming, A. M.; Zhou, J.; Wallace, S. S.; Burrows, C. J., A role for the fifth G-track in G-quadruplex forming oncogene promoter sequences during oxidative stress: Do these “spare tires” have an evolved function? ACS Cent. Sci. 2015, 1, 226–233.

17. Burra, S.; Marasco, D.; Malfatti, M. C.; Antoniali, G.; Virgilio, A.; Esposito, V.; Demple, B.; Galeone, A.; Tell, G., Human AP-endonuclease (Ape1) activity on telomeric G4 structures is modulated by acetylatable lysine residues in the N-terminal sequence. DNA Repair 2019, 73, 129–143.

18. Zhang, A. Y.; Balasubramanian, S., The kinetics and folding pathways of intramolecular G-quadruplex nucleic acids. J. Am. Chem. Soc. 2012, 134, 19297–19308.

19. Gray, R. D.; Chaires, J. B., Kinetics and mechanism of K+- and Na+-induced folding of models of human telomeric DNA into G-quadruplex structures. Nucleic Acids Res. 2008, 36, 4191–4203.

20. Crossley, M. P.; Bocek, M.; Cimprich, K. A., R-Loops as cellular regulators and genomic threats. Mol. Cell 2019, 73, 398–411.

21. McNeill, D. R.; Whitaker, A. M.; Stark, W. J.; Illuzzi, J. L.; McKinnon, P. J.; Freudenthal, B. D.; Wilson, D. M., 3rd, Functions of the major abasic endonuclease (APE1) in cell viability and genotoxin resistance. Mutagenesis 2020, 35, 27–38.

22. Pan, L.; Zhu, B.; Hao, W.; Zeng, X.; Vlahopoulos, S. A.; Hazra, T. K.; Hegde, M. L.; Radak, Z.; Bacsi, A.; Brasier, A. R.; Ba, X.; Boldogh, I., Oxidized guanine base lesions function in 8-oxoguanine DNA glycosylase1-mediated epigenetic regulation of nuclear factor kappaB-driven gene expression. J. Biol. Chem. 2016, 291, 25553–25566.

23. Robinson, J.; Raguseo, F.; Nuccio, S. P.; Liano, D.; Di Antonio, M., DNA G-quadruplex structures: more than simple roadblocks to transcription? Nucleic Acids Res. 2021, 49, 8419–8431.

24. Williams, S. L.; Casas-Delucchi, C. S.; Raguseo, F.; Guneri, D.; Li, Y.; Minamino, M.; Fletcher, E. E.; Yeeles, J. T.; Keyser, U. F.; Waller, Z. A.; Di Antonio, M.; Coster, G., Replication-induced DNA secondary structures drive fork uncoupling and breakage. EMBO J. 2023, 42, e114334.

25. Spiegel, J.; Cuesta, S. M.; Adhikari, S.; Hänsel-Hertsch, R.; Tannahill, D.; Balasubramanian, S., G-quadruplexes are transcription factor binding hubs in human chromatin. Genome Biol. 2021, 22, 117.

26. Irving, K. L.; King, J. J.; Waller, Z. A. E.; Evans, C. W.; Smith, N. M., Stability and context of intercalated motifs (i-motifs) for biological applications. Biochimie 2022, 198, 33–47.

27. Cheng, M.; Qiu, D.; Tamon, L.; Ištvánková, E.; Víšková, P.; Amrane, S.; Guédin, A.; Chen, J.; Lacroix, L.; Ju, H.; Trantírek, L.; Sahakyan, A. B.; Zhou, J.; Mergny, J. L., Thermal and pH stabilities of i-DNA: Confronting in vitro experiments with models and in-cell NMR data. Angew. Chem. Int. Ed. Engl. 2021, 60, 10286–10294.

