## Supplemental Information for "DNA Damage Accelerates G-Quadruplex Folding in a Duplex-G-Quadruplex-Duplex Context"

Aaron M. Fleming,* Brandon Leonel Guerra Castañaza Jenkins,

Bethany A. Buck,* Cynthia J. Burrows*

Department of Chemistry, University of Utah, 315 S. 1400 East, Salt Lake City, UT

84112-0850 United States

**Item	Page**

**Methods**	S2

**Figure S1**. DNA strand purity and ESI-MS analysis.	S7

**Figure S2**. Comparison of the *VEGF*-DGD ^1^H-NMR and CD data to the

*cMYC*-DGD reported.	S9

**Figure S3**. Thermal melting analysis of the *VEGF*-DGD construct	S10

**Figure S4**. Observed rate constants measured to calculate folding half-lives.	S11

**Figure S5.** Folding rates for *VEGF*-DGD motifs with variable length poly-T

complements.	S12

**Figure S6.** The CD and NMR spectra recorded before and after isothermal

folding of the PQSs.	S13

**Figure S7.** Fluorescence anisotropy analysis measuring the APE1:

*VEGF*-DGD binding interaction dissociation constants.	S14

**Figure S8.** The CD spectra before and after folding with APE1 present.	S15

**Figure S9.** The CD spectra before and after folding with DNA damage

in the 5′-G or 3′-G runs.	S16

**References**	S18

**Methods**

***DNA and Protein Preparation***

All DNA oligomers were synthesized by the DNA/Peptide core facility at the University of Utah to obtain the 5′-detritylated DNA strands. Site-specific introduction of OG or the abasic site analog tetrahydrofuran (F) was achieved using commercially available phosphoramidites (Glen Research). As described in Figure S1, we found the standard deprotection method to be inadequate to obtain the purest DNA possible free of sequences that retained protecting groups and/or cyanoethylation of the T nucleotide.^1^ Two different cleavage and deprotection approaches were used during this study. In the first method, DNA oligomers were cleaved from the synthesis column in aqueous 30% NH_4_OH and 200 mM BME at room temperature for 24 h, followed by deprotection in the same solvent system by heating the samples at 55 °C for 24 h. In the second method, DNA oligomers were cleaved and deprotected in the gas-phase with 25 mL each of aqueous 40% methylamine and aqueous 30% NH_4_OH at 85 °C for 1 h. Using either cleavage and deprotection method, the crude samples were lyophilized to dryness and then anion-exchange HPLC purified.

The crude oligomers were purified using a semi-preparative anion-exchange HPLC column running a mobile phase system consisting of A (1:9 MeCN:ddH_2_O water) and B (1 M LiCl, 20 mM LiOAc at pH 7 in 1:9 MeCN:ddH_2_O). The method was initiated at 1% B and increased to 100% B via a linear gradient over 30 min with a flow rate of 3 mL/min while monitoring the absorbance at 260 nm. Purified samples were dialyzed against ddH_2_O for 36 h to remove purification salts, then lyophilized to dryness and resuspended in ddH_2_O. Sample concentrations were determined by measuring the absorbance at 260 nm and using the nearest neighbor approximation to obtain extinction coefficients for calculating purified stock DNA sample concentrations. Extinction coefficients for the F-containing DNA strands were estimated by omitting a nucleotide for the site and the OG-containing strands used G at this site in the calculations. Purity of the oligomers was determined using an analytical anion-exchange HPLC column running a mobile phase system consisting of A (1:9 MeCN:ddH_2_O water) and B (1 M LiCl, 20 mM LiOAc at pH 7 in 1:9 MeCN:ddH_2_O). The method was initiated at either 15 or 25% B and increased to 100% B via a linear gradient over 30 min with a flow rate of 1 mL/min while monitoring the absorbance at 260 nm.

Thermodynamic folding of the DGD constructs was achieved by heating the G-rich strand (50 μM) to 90 °C in a water bath for 5 min in a buffer system comprised of 20 mM KP_i_ (pH 6.8) and 120 mM KCl. After the thermal denaturation, the heat was turned off and the water bath was allowed to reach room temperature (~3 h). Next, the complementary strand with the poly-T tether (final concentration = 60 μM) was added to the G4 annealed G-rich strand, and then the mixture was placed at 4 °C for 24 h to allow the duplex arms to fold before analysis. This annealing approach follows the prior report on the *c-MYC*-DGD motif.^2^

Isothermal folding of the DGD systems was achieved by first mixing the G-rich strand (50 μM) with the complementary strand (60 μM) in a buffer system comprised of 20 mM Tris (pH 7.2) and 140 mM LiCl. The mixture was placed at 4 °C for 24 h before further folding, which is described below.

The APE1 protein used in the present studies had a D210A mutation to eliminate the catalytic endonuclease activity while retaining the ability to bind DNA. The protein was prepared as previously described by our laboratory.^3^

***Circular Dichroism (CD) Analysis***

For CD analysis, the thermodynamically folded DNA samples at a 10 μM concentration were placed in a 0.2-cm quartz cuvette at 20 °C. The CD spectra were recorded from 200-350 nm on a Jasco J815 system. The spectra for the DNA strands had the buffer background subtracted and then the ellipticity values obtained were converted to molar ellipticity values for visualization by plotting [θ] on the y-axis and wavelength (nm) on the x-axis.

For isothermal folding experiments, the DNA to be analyzed was placed in a 1.0-cm quartz cuvette held at 30 °C via the Peltier heating module connected to the CD spectrometer. Samples were rapidly mixed with a stir bar controlled via the spectrometer. After thermal equilibration for 10 min, a bolus of one of three different folding reagent solutions was added to the mixing sample while monitoring the CD signal at 260 nm every second for the samples that folded on the minutes time scale, or 50 msec for those that folded on the seconds timescale: (1) A KCl solution was added to achieve a final 1 μM DNA concentration and 100 mM KCl concentration.; (2) A KCl and MgCl_2_ mixture was added to achieve final concentrations of 1 μM DNA, 100 mM KCl, and 10 mM MgCl_2_; (3) A KCl, MgCl_2_, and D210A-APE1 mixture was added to achieve final concentrations of 1 μM DNA, 1 μM D210A-APE1, 100 mM KCl, and 10 mM MgCl_2_. The folding process was monitored for up to 30 min. Time-dependent CD data at 260 nm were fit to a first-order exponential function as previously described to obtain the observed rate constants (Eq. 1).^4^

$y\left( t \right)=yo+Ae^{(-\frac{t}{t1})}$	Eq. 1

where y(t) is the raw CD signal at time *t* and yo, A, and *t_1_* are fitting parameters. *t_1_* is the characteristic folding time, in which 1/*t_1_* is the rate reported in this work. Folding half-live values were obtained from the observed rate constants measured. It must be noted that previous kinetic rate measurements for PQSs folding to G4 motifs found this to be a multistep process, though these measurements were conducted on PQS systems folding to G4s outside the influence of duplex DNA anchors as used in the present system.^5,6^ Attempts to fit the various *VEGF*-DGD construct kinetic rate data to a bi-exponential function resulted in less favorable fitting statistics. The present work is focused on whether the PQS can fold when restrained by flanking duplex regions and a modeled complementary strand, as opposed to attempting to delineate the folding pathway details. Therefore, we have chosen the single-exponential fitting equation for the comparative analysis presented. For each sample, the CD spectrum from 200-350 nm was recorded before and after the folding reaction and processed as previously described.

***1D* *^1^H-NMR Analysis***

All 1D ^1^H-NMR spectra of the thermodynamically and isothermally prepared *VEGF*-DGD construct samples were collected using a WATERGATE (WATER-suppression by GrAdient-Tailored Excitation) water suppression pulse sequence^7,8^ at 298 K on a Varian VNMRS 800-MHz spectrometer. Spectra were collected at DNA concentrations ranging from 25 μM-100 μM using a respective number of 512-2048 scans. For analysis of the various thermodynamically folded *VEGF*-DGD constructs (prepared as described above), a buffering system of 20 mM KP_i_ (pH 6.8), 120 mM KCl, and 10% D_2_O was utilized. Analysis for the various isothermally folded *VEGF*-DGD constructs was a two-fold process. First, duplex formation within the various *VEGF*-DGD systems was achieved as described above, and NMR measurements in a buffering system of 20 mM Tris (pH 6.8), 140 mM LiCl, and 10% D_2_O was performed. Second. A final concentration of 100 mM KCl was added to each Li^+^ samples, allowed to incubate for 1 h at room temperature, prior to spectral acquisition. All data were processed using NMRPipe,^9^ and analyzed using the nmrDraw module.

***Thermal Melting (T_M_) Analysis***

The unfolded-DGD system in LiCl and buffer for thermal melting (*T_m_*) analysis was prepared as described above and then diluted to 1 μM DNA concentration. The samples were placed in a quartz *T_m_* analysis cuvette that was placed in a temperature-regulated UV-vis spectrometer (Shimadzu UV-1800) followed by thermal equilibration at 20 °C for 5 min before the commencement of the experiment. The thermally induced denaturation of the G4s was monitored at 295 nm by heating the sample from 20 to 100 °C at a ramp rate of 0.5 °C/min followed by a 60-sec equilibration and then measuring the absorbance value at 295 nm. The thermal induced denaturation of the duplex handles was monitored by the absorbance value at 260 nm. The absorbance data collected were background subtracted, and then *T_m_* values were determined using Shimadzu’s *T_m_* analysis software.^10^

***Fluorescence anisotropy binding assays***

The fluorescence anisotropy experiments were conducted by titrating the prepared 5′-FAM-labeled DNA samples with the appropriate APE1 protein from 0-5000 nM followed by incubating the mixture at 22 °C for 30 min before analysis. The fluorescence anisotropy measurements were carried out on a BioTek Synergy2 Multi-Mode Microplate Reader at excitation and emission wavelengths 485 and 520 nm, respectively. The anisotropy values (r) were calculated with Eq. 2 in which *I_par_* is the parallel emission intensity and *I_per_* is the perpendicular emission intensity.

$r=\frac{I_{par}-I_{per}}{I_{par}+{2*I}_{per}}$	Eq. 2

The r values obtained were plotted against the log[APE1] to produce sigmoidal curves that were fit to the following Hill equation where the bottom is the lower plateau value of r and top is the higher plateau value of r defining the sigmoidal curve. The dissociation-binding constant is *K_D_* and *n* is the Hill coefficient (Eq. 3).^11^

$r=bottom+\frac{\left( top-bottom \right)}{1+{10^\left( \left. logKd-log[APE1] \right) \right.}^{n}}$	Eq. 3

The *K_D_* and n values were determined from the equation obtained that best fit the data points based on linear least-squares analysis. The error bars in the values reported are the standard deviations of the values obtained from triplicate trials.

**Data Availability**

All data are available upon request.

**Figure S1**. DNA strand purity and ESI-MS analysis.

A known issue with synthesis of long DNA, and sequences rich in either T or G nucleotides is the presence of impurities after their synthesis. These problematic sequences represent all of the DNA oligomers studied herein. The most common impurities include cyanoethylation at N3 of the T nucleotide or failure to release the exocyclic amine protecting groups from A, C, and most significantly for G. These impurities can be observed when analyzing the DNA with analytical anion-exchange HPLC, as we have used in our work. When we commenced this study, we found the DNA cleaved and deprotected under standard conditions led to high levels of impurities (data not shown). Mitigation of this issue was achieved by three changes to the synthesis of the DNA strands studied. (1) The more labile dG phosphoramidite with an *N^2^*-dmf protecting group was used for synthesis. (2) The cleavage and deprotection conditions were modified to run for longer times to drive release of the protecting groups. (3) Addition of a trapping nucleophile was introduced to prevent cyanoethylation of the T nucleotides. Two different approaches were employed to have longer deprotecting times with a trapping nucleophile present (see methods section for complete details of these changes). Lastly, all DNA were HPLC purified before study. Below are example analytical anion-exchange HPLC chromatograms demonstrating the purities of the DNA oligomers are >90% (panels A-D); and we conducted ESI^-^-MS to verify the purities for two of the longer strands (panels E and F).

**Example Analytical Anion-exchange HPLC Chromatograms**

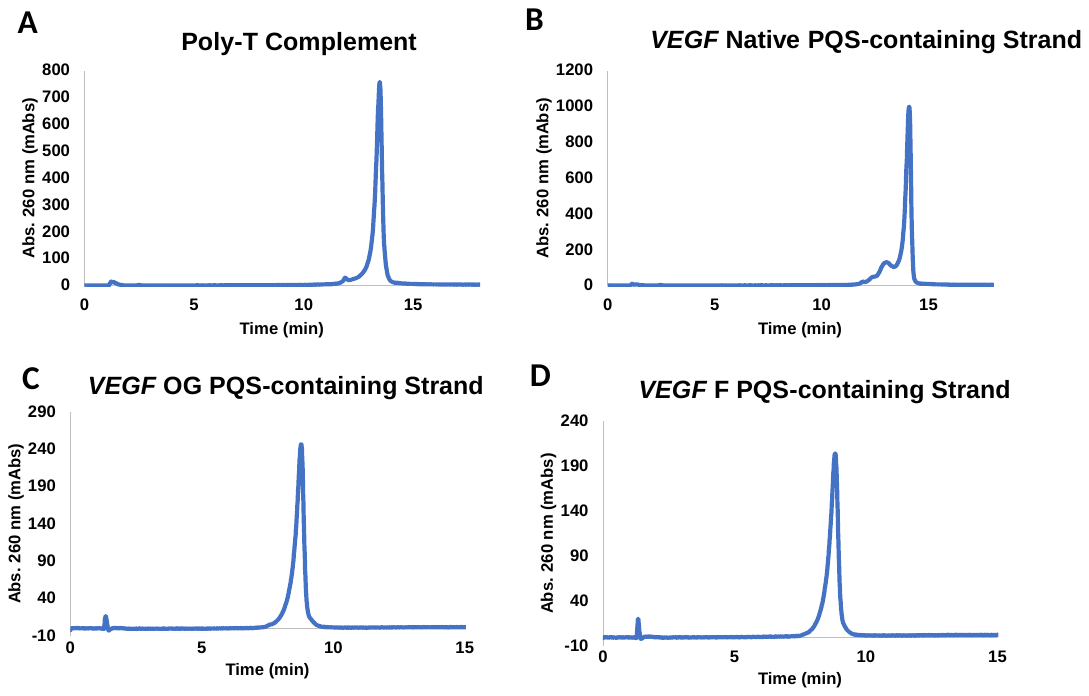

**ESI^-^-MS Spectra**

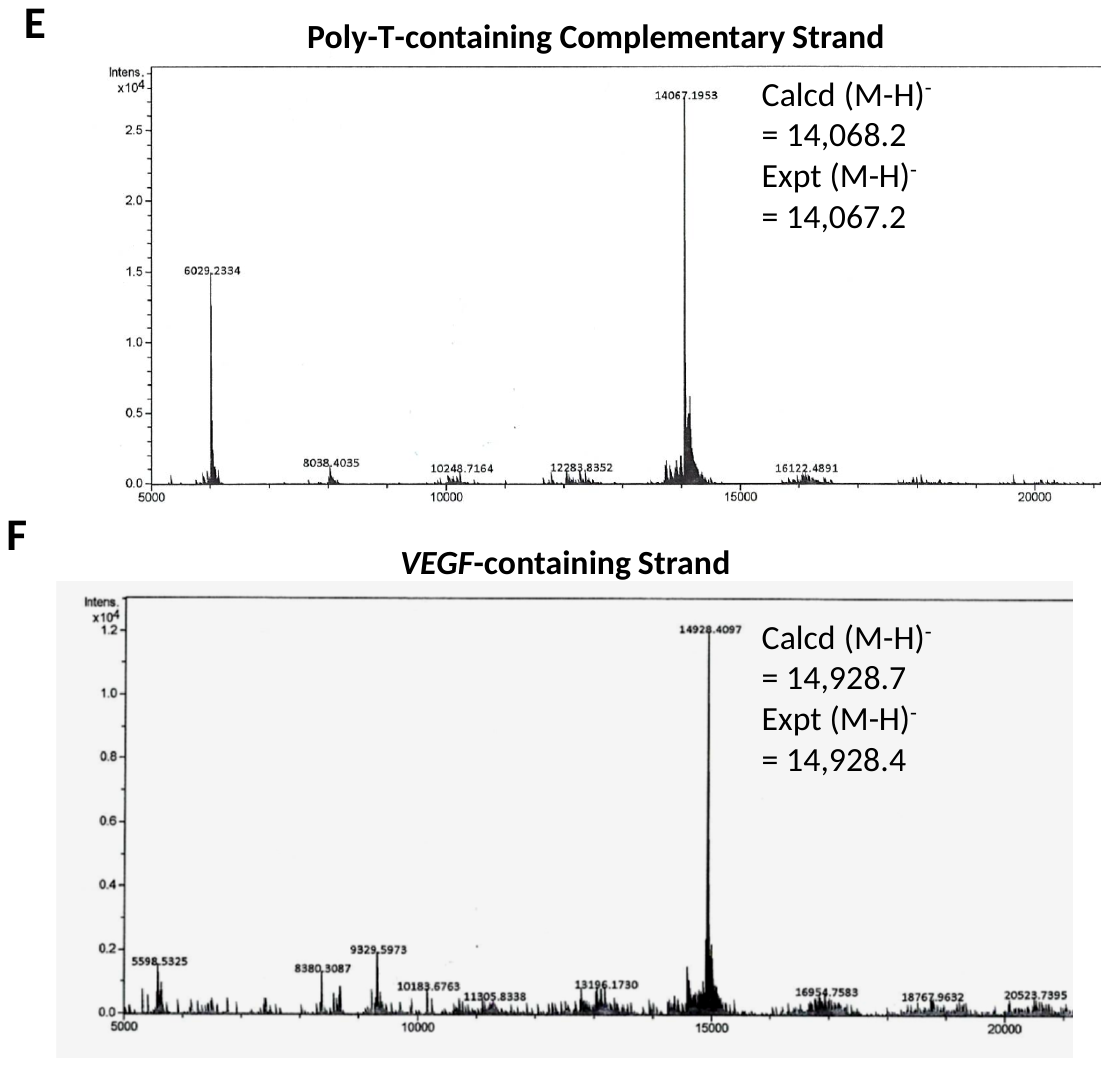

**Figure S2**. Comparison of the *VEGF*-DGD ^1^H-NMR and CD data to the *cMYC*-DGD reported. The data for the *c-MYC*-DGD construct was previously reported.^2^

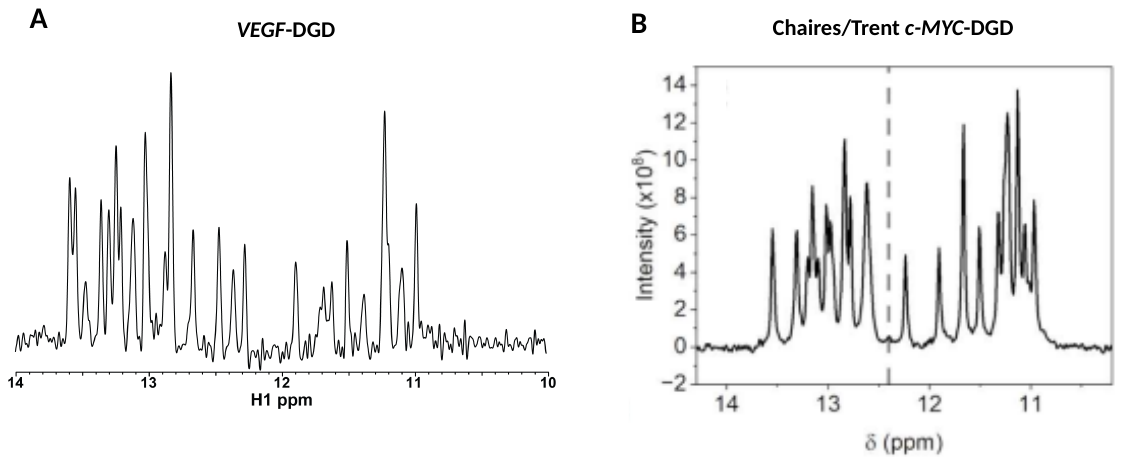

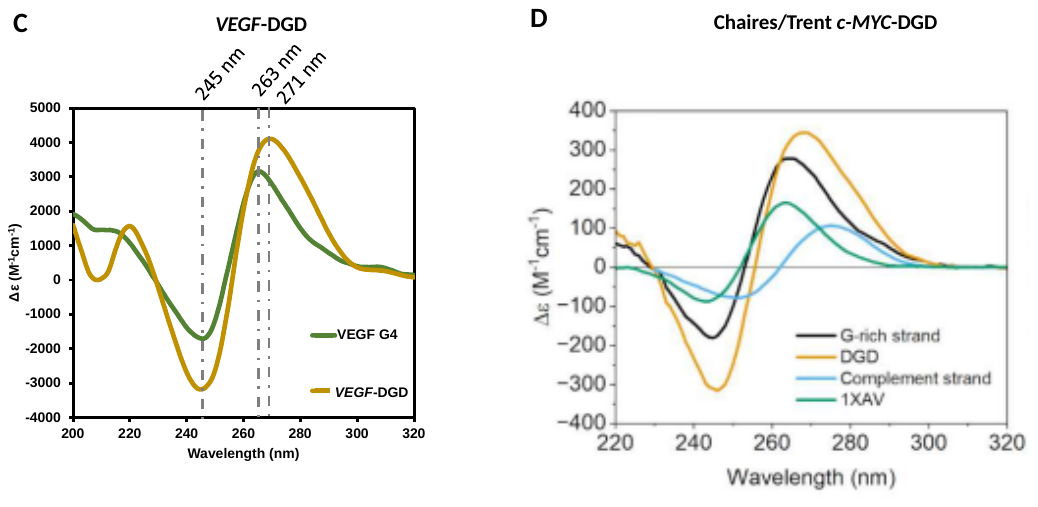

Comparison of 1D ^1^H-NMR spectra for (A) *VEGF*-DGD and (B) *cMYC*-DGD constructs. The *VEGF*-DGD sample was prepared under thermodynamic folding conditions in 20 mM KP_i_ (pH 6.8) and 120 mM KCl followed by analysis at 25 °C on an 800-MHz NMR. The *cMYC*-DGD spectrum from the literature^2^ was obtained on a sample analyzed in 8 mM NaP_i_ (pH 7.2) and 185 mM KCl on a 600-MHz NMR at 20 °C. Comparison of the thermodynamically folded CD spectra for (C) *VEGF*-DGD to the (D) *cMYC*-DGD. The comparisons are for the G4s folded outside the DGD context shown with the green lines, and the G4s in the DGD context shown with the yellow lines. The CD samples were both analyzed at 20 °C in the same buffer and salt as employed in the NMR experiments.

**Figure S3**. Thermal melting analysis of the *VEGF*-DGD construct.

The experimental T_m_ curve for 1 µM of the isothermally prepared *VEGF*-DGD (buffer conditions: 20 mM Tris (pH 7.2), 140 mM LiCl) was biphasic with inflection points at 39 °C and 52 °C.

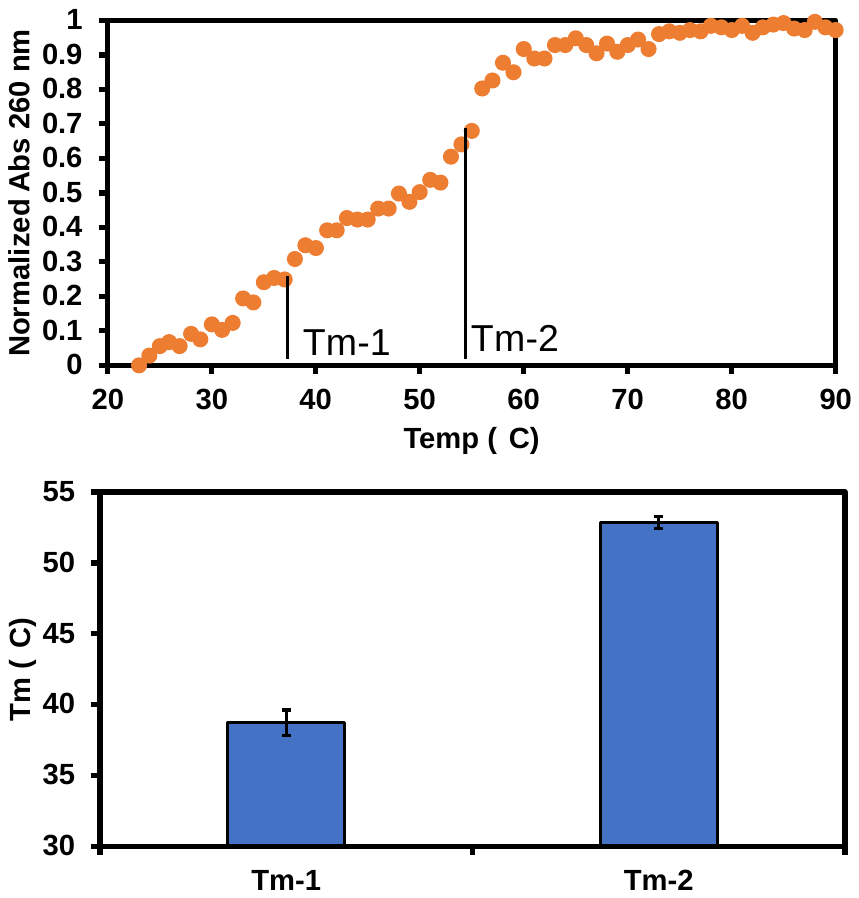

**Figure S4**. Observed rate constants measured to calculate folding half-lives.

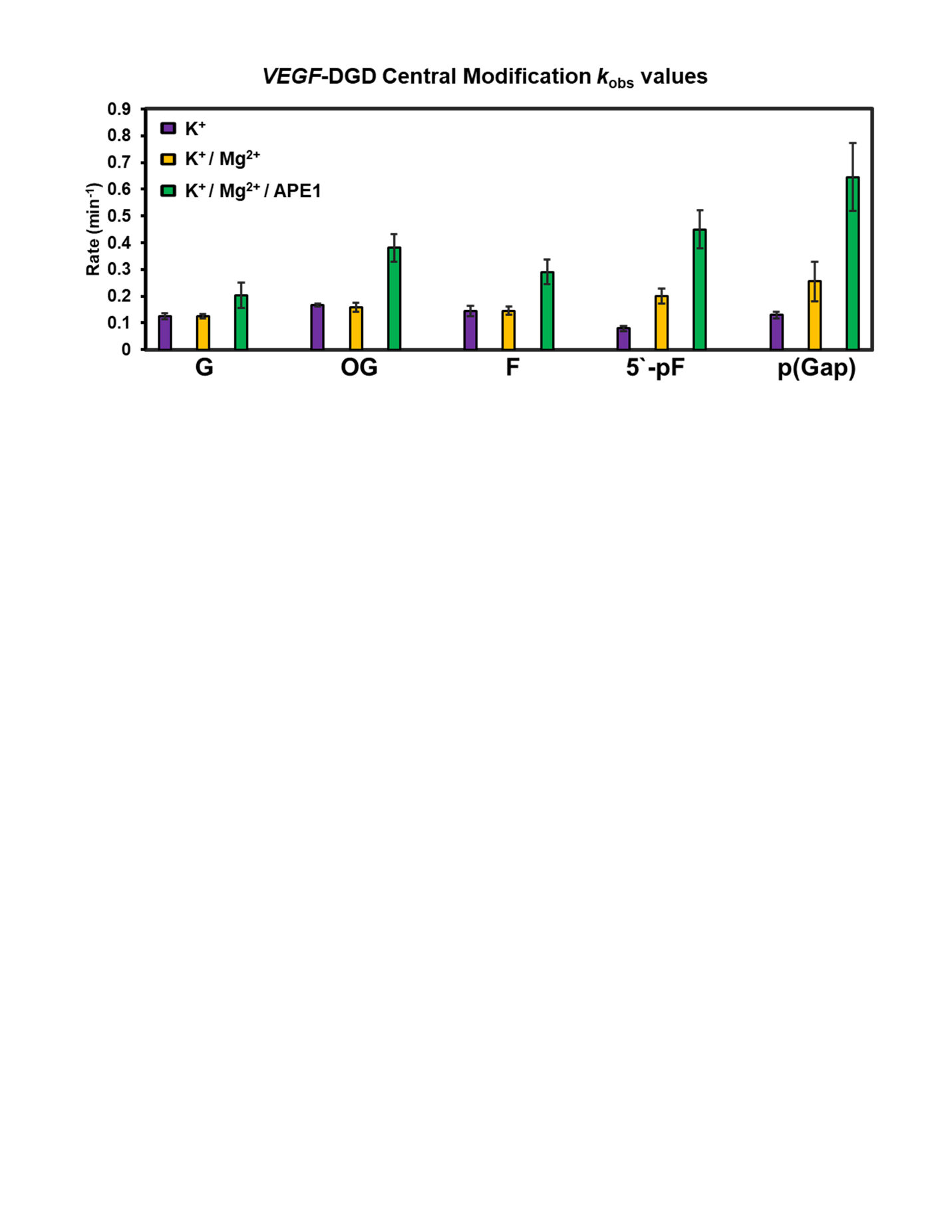

**Figure S5.** Folding rates for *VEGF*-DGD motifs with variable length poly-T complements.

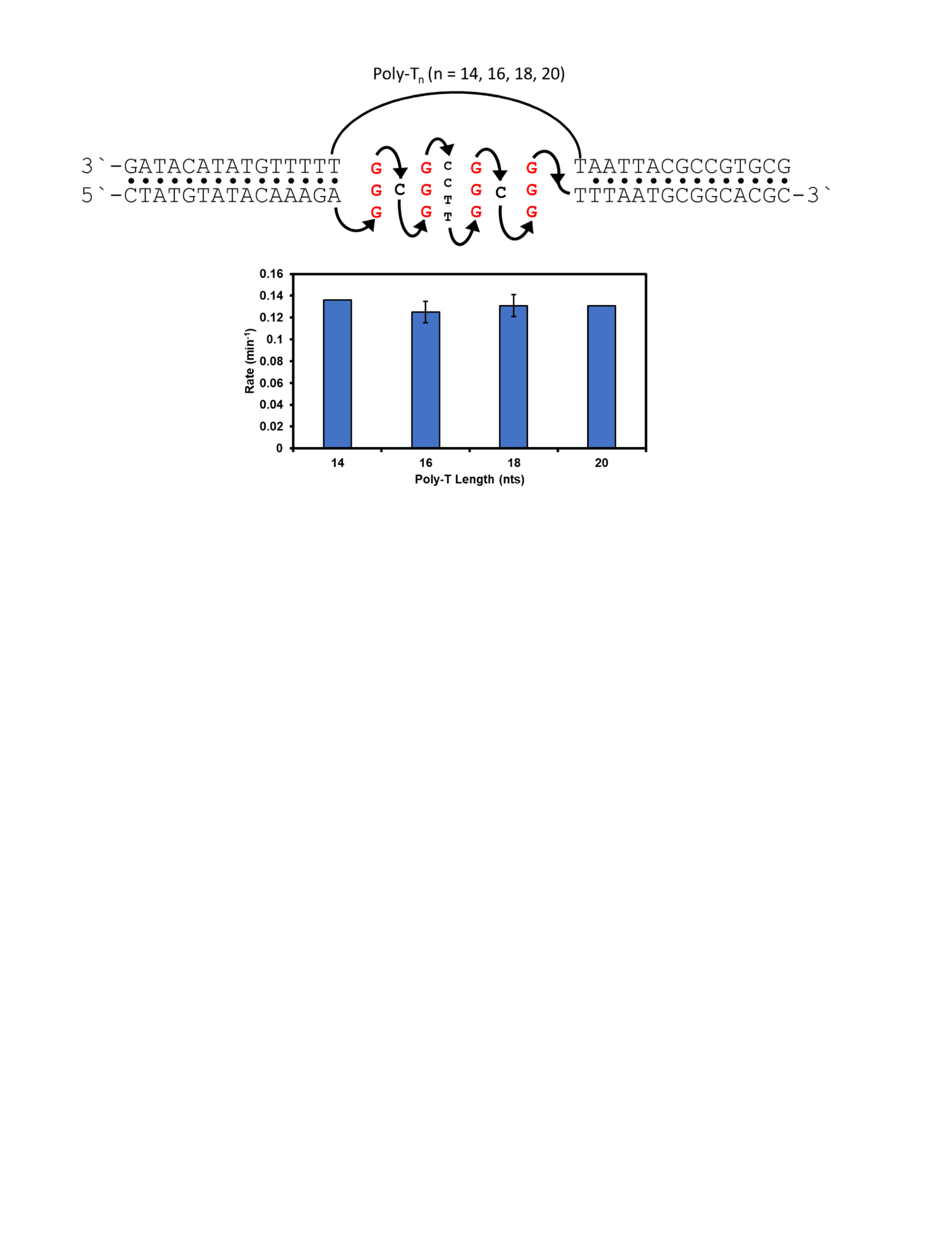

**Figure S6.** CD and 1D ^1^H-NMR spectra recorded before and after isothermal folding of the PQSs.

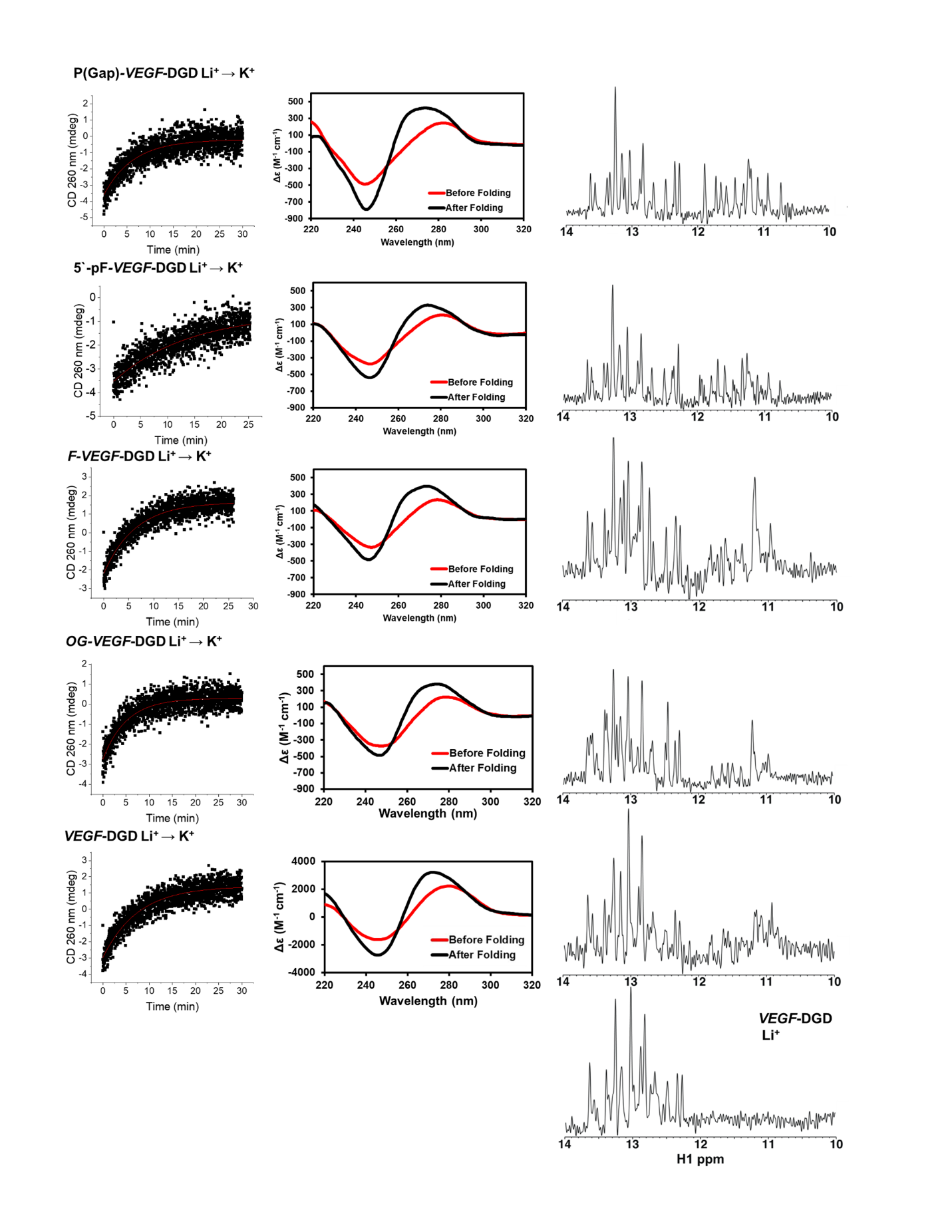

**Figure S7.** Fluorescence anisotropy analysis measuring the APE1:*VEGF*-DGD binding interaction dissociation constants.

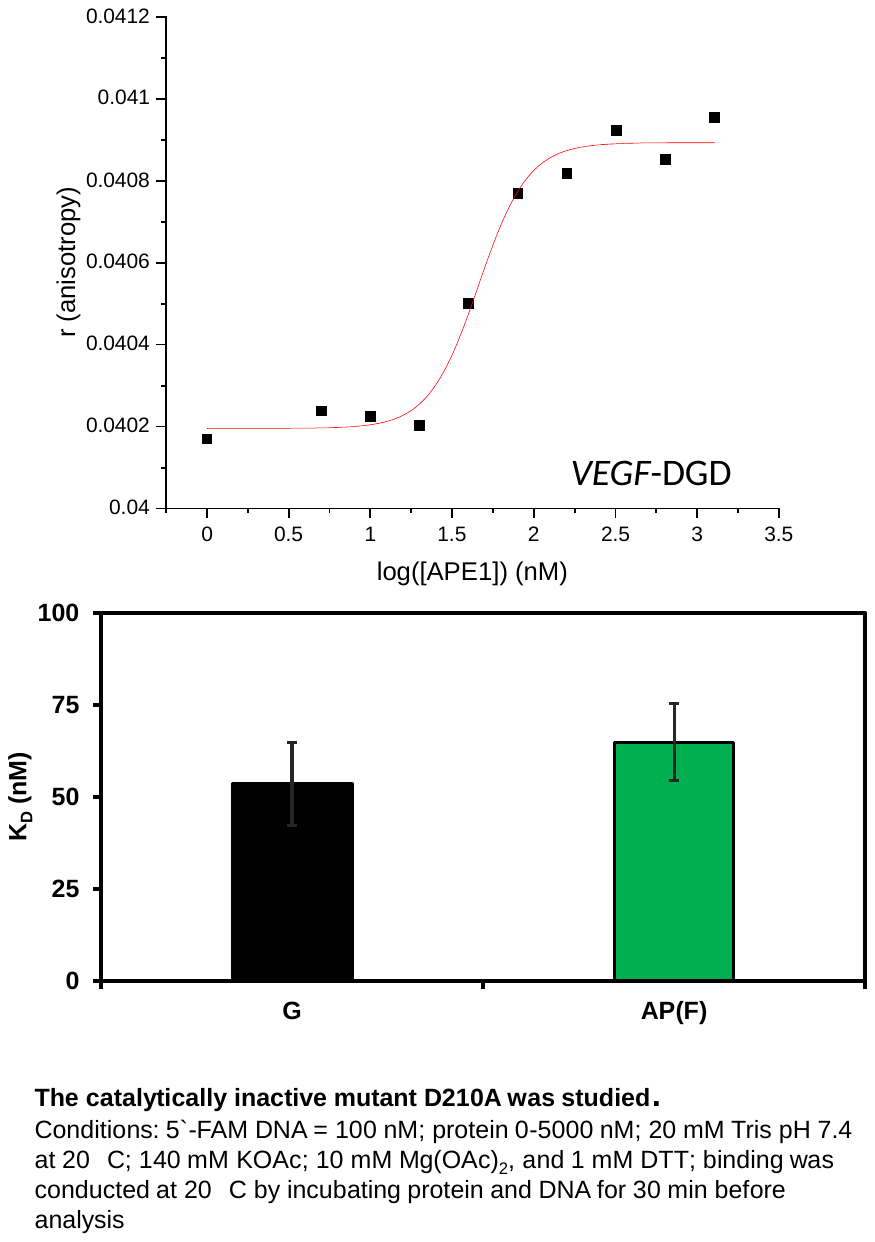

Conditions: 5`-FAM DNA = 100 nM; protein 0-5000 nM; 20 mM Tris pH 7.4 at 20 °C; 140 mM KOAc; 10 mM Mg(OAc)_2_, and 1 mM DTT; binding was conducted at 20 °C by incubating protein and DNA for 30 min before analysis. The D210A-APE1 was used for the binding assay as we previously described.^3^

**Figure S8.** CD spectral analysis for folding of various *VEGF*-DGD constructs before and after folding with APE1 present.

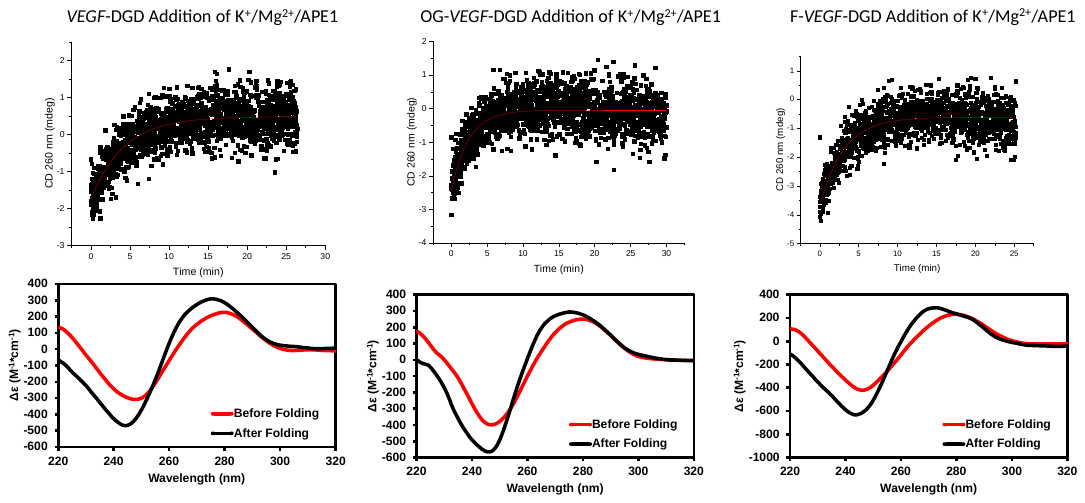

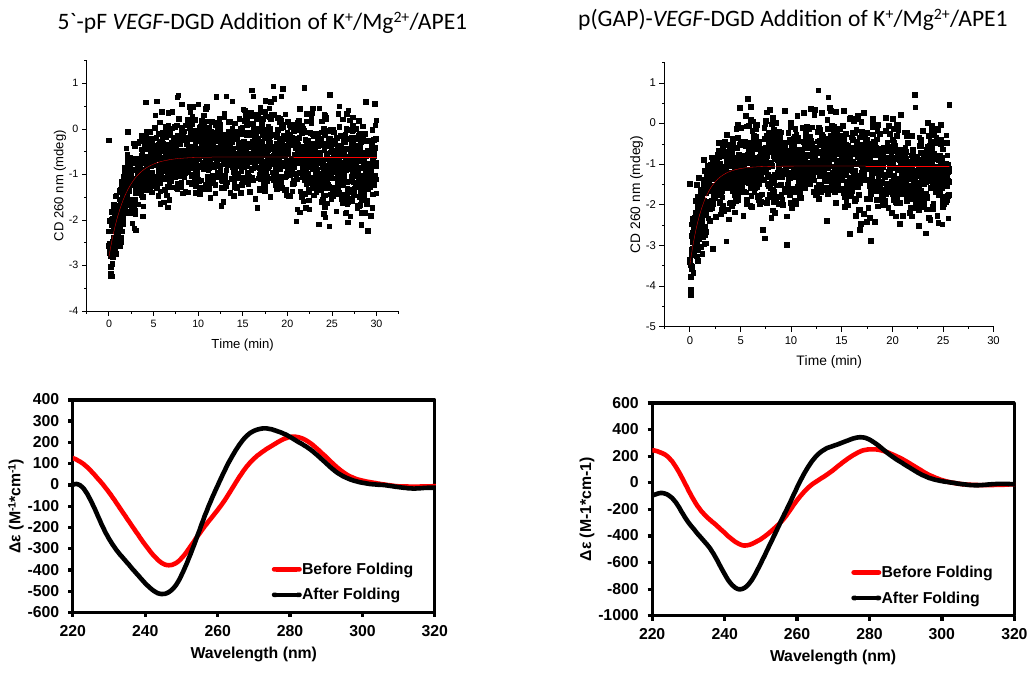

**Figure S9.** CD spectral analysis for folding of *VEGF*-DGD constructs with DNA damage in the 5′-G or 3′-G runs. The time-dependent CD at 260 nm and CD spectra before and after the addition are plotted for addition of K^+^. The position-dependent data VEGF-DGD constructs are shown for (A) 5`-G run OG, (B) 5`-G run F, (C) 5`-G run 5`-pF, (D) 5`-G run p(Gap), (E) 3`-G run OG, (F) 3`-G run F, (G) 3`-G run 5`-pF, and (H) 3`-G run p(Gap). For the DGD systems that folded in minutes, the time-dependent CD data were collected every 1 sec, and for those that folded in the seconds, the CD were recorded either every 50 msec.

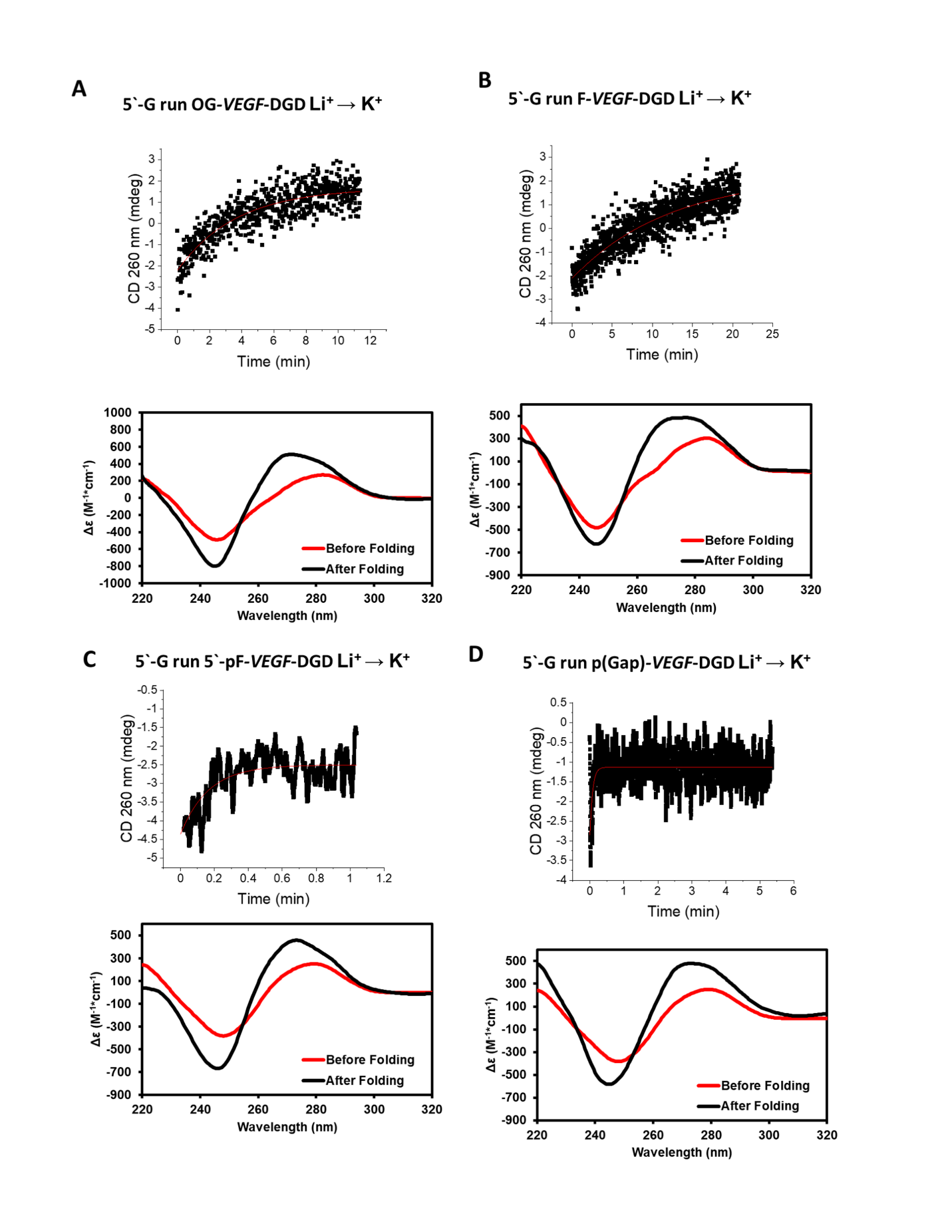

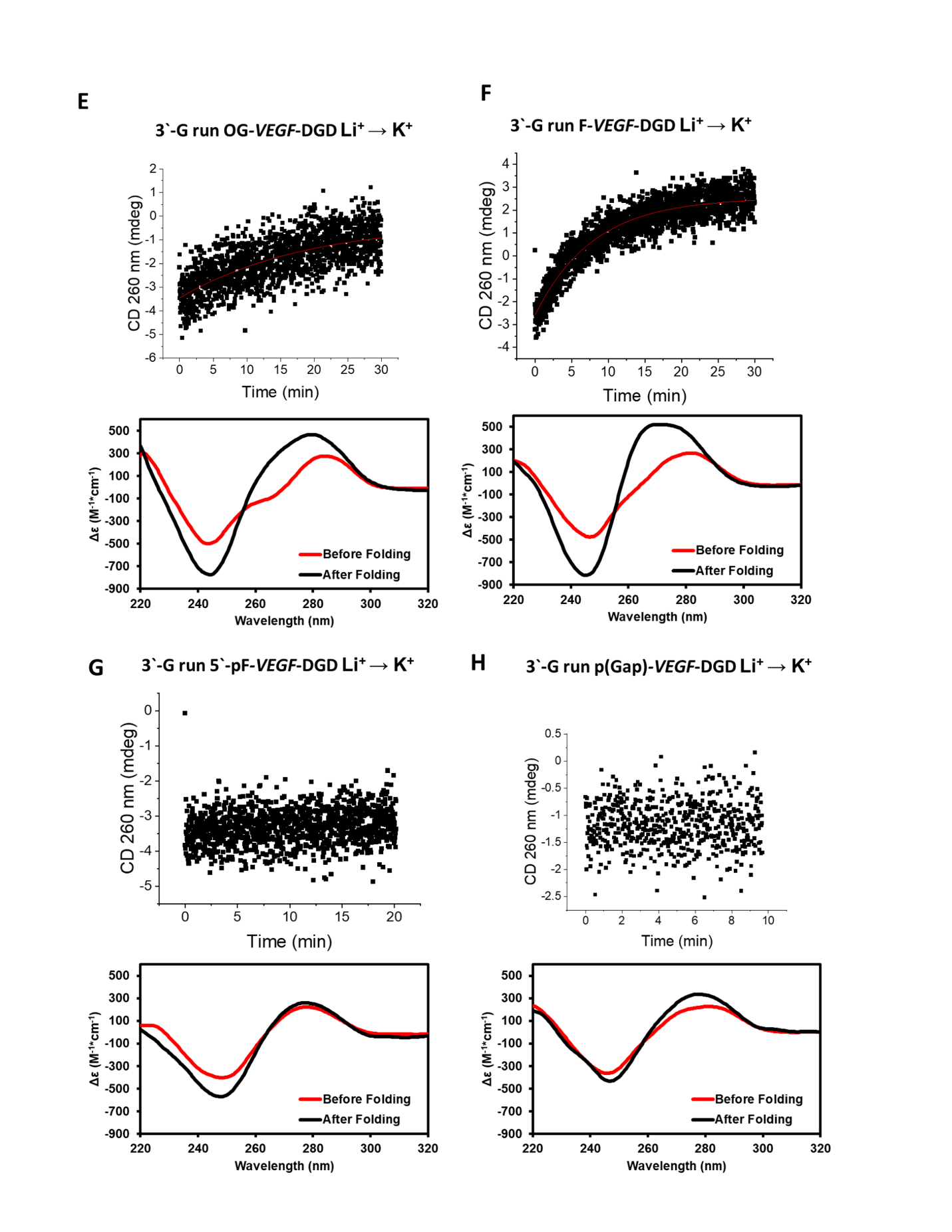

**References**

1.	Tsunoda, H.; Kudo, T.; Ohkubo, A.; Seio, K.; Sekine, M., Synthesis of oligodeoxynucleotides using fully protected deoxynucleoside 3'-phosphoramidite building blocks and base recognition of oligodeoxynucleotides incorporating N3-cyano-ethylthymine. *Molecules (Basel, Switzerland)* **2010,** *15*, 7509-7531.

2.	Monsen, R. C.; Chua, E. Y. D.; Hopkins, J. B.; Chaires, J. B.; Trent, J. O., Structure of a 28.5 kDa duplex-embedded G-quadruplex system resolved to 7.4 Å resolution with cryo-EM. *Nucleic Acids Res.* **2023,** *51*, 1943-1959.

3.	Fleming, A. M.; Howpay Manage, S. A.; Burrows, C. J., Binding of AP endonuclease-1 to G-quadruplex DNA depends on the N-terminal domain, Mg^2+^, and ionic strength. *ACS Bio. & Med. Chem. Au* **2021,** *1*, 44-56.

4.	Chen, C.; Li, M.; Xing, Y.; Li, Y.; Joedecke, C. C.; Jin, J.; Yang, Z.; Liu, D., Study of pH-induced folding and unfolding kinetics of the DNA i-motif by stopped-flow circular dichroism. *Langmuir* **2012,** *28*, 17743-17748.

5.	Gray, R. D.; Chaires, J. B., Kinetics and mechanism of K^+^- and Na^+^-induced folding of models of human telomeric DNA into G-quadruplex structures. *Nucleic Acids Res.* **2008,** *36*, 4191-4203.

6.	Zhang, A. Y.; Balasubramanian, S., The kinetics and folding pathways of intramolecular G-quadruplex nucleic acids. *J. Am. Chem. Soc.* **2012,** *134*, 19297-19308.

7.	Piotto, M.; Saudek, V.; Sklenar, V., Gradient-tailored excitation for single-quantum NMR spectroscopy of aqueous solutions. *J. Biomol. NMR* **1992,** *2*, 661-665.

8.	Liu, M.; Mao, X.-a.; Ye, C.; Huang, H.; Nicholson, J. K.; Lindon, J. C., Improved WATERGATE Pulse Sequences for Solvent Suppression in NMR Spectroscopy. *J. Magn. Res.* **1998,** *132*, 125-129.

9.	Delaglio, F.; Grzesiek, S.; Vuister, G. W.; Zhu, G.; Pfeifer, J.; Bax, A., NMRPipe: a multidimensional spectral processing system based on UNIX pipes. *J. Biomol. NMR* **1995,** *6*, 277-293.

10.	Mergny, J. L.; Lacroix, L., Analysis of thermal melting curves. *Oligonucleotides* **2003,** *13*, 515-537.

11.	Gesztelyi, R.; Zsuga, J.; Kemeny-Beke, A.; Varga, B.; Juhasz, B.; Tosaki, A., The Hill equation and the origin of quantitative pharmacology. *Arch. Hist. Exact. Sci.* **2012,** *66*, 427-438.
